# Structure of the full-length VEGFR-1 extracellular domain in complex with VEGF-A

**DOI:** 10.1101/102822

**Authors:** Sandra Markovic-Mueller, Edward Stuttfeld, Mayanka Asthana, Tobias Weinert, Spencer Bliven, Kenneth N. Goldie, Kaisa Kisko, Guido Capitani, Kurt Ballmer-Hofer

**Affiliations:** Paul Scherrer Institute, Laboratory of Biomolecular Research, 5232 Villigen PSI, Switzerland; Biozentrum, University of Basel, Basel, Switzerland; Center for Cellular Imaging and Nano Analytics (C-CINA), Biozentrum, University of Basel, Basel, Switzerland

## Abstract

Vascular Endothelial Growth Factors (VEGFs) regulate blood and lymph vessel development upon activation of three receptor tyrosine kinases (RTKs), VEGFR-1, −2, and −3. Partial structures of VEGFR/VEGF complexes based on single particle electron microscopy, small angle X-ray scattering, and X-ray crystallography revealed the location of VEGF binding and domain arrangement of individual receptor subdomains. Here we describe the structure of the full-length VEGFR-1 extracellular domain (ECD) in complex with VEGF-A at 4 Å resolution. We combined X-ray crystallography, single particle electron microscopy, and molecular modeling for structure determination and validation. The structure reveals the molecular details of ligand-induced receptor dimerization, in particular of homotypic receptor interactions in Ig-domains 4, 5, and 7. Functional analyses of ligand binding and receptor activation confirm the relevance of these homotypic contacts and identify them as potential therapeutic sites to allosterically inhibit VEGFR-1 activity.

## INTRODUCTION

The formation of functional blood and lymphatic vessels is a keystone during embryo development and is essential for supplying all organs with oxygen and nutrients and for disposal of catabolites. Vascular Endothelial Growth Factors (VEGFs) are the main drivers of vasculogenesis, the *de novo* formation of vessels, and angiogenesis, the formation of new vessels from preexisting vasculature (reviewed in (Smith et al., 2015; Shibuya, 2014; Moens et al., 2014). VEGFs activate three VEGF receptors (VEGFR-1, −2 and −3), representing the type V subfamily of receptor tyrosine kinases (RTKs) in the human kinome and are indispensable for vessel development. The molecular details of the regulatory processes specific for this RTK subfamily are therefore of high medical relevance.

Mutation or ablation of a specific VEGFR gave rise to distinct disease profiles (Shibuya, 2014) documenting signal diversity among these three receptors. Both ablation of VEGFR-1 and −2 were embryonic lethal; the former due to complete disorganization of blood vessels (Fong et al., 1995), the latter because of the complete absence of endothelial cells (Shalaby et al., 1995). Interestingly, expression of a kinase deficient VEGFR-1 variant did not lead to embryonic lethality and mice expressing this receptor variant had only minor defects (Hiratsuka et al., 1998; Hiratsuka et al., 2001). Deregulated expression of a soluble splice variant of VEGFR-1, encompassing only the extracellular domain (ECD) and acting as a ligand trap(Kendall and Thomas, 1993), is associated with vascular defects such as the placental deficiency preeclampsia (Luttun and Carmeliet, 2003; Maynard et al., 2003), or corneal and retinal avascularity (Luo et al., 2013; Ambati et al., 2006). This variant is thus a potentially interesting target for drug development.

RTK activation requires ligand-mediated dimer or multimer formation, which instigates transmembrane signaling resulting in activation of the intracellular kinase domain (Lemmon and Schlessinger, 2010; Lemmon and Schlessinger, 1994). The exact structural changes driving this process are unique for each RTK subfamily and are not known for VEGF receptors. Up to now only limited structural information is available for VEGFRs. A low resolution structure of a VEGFR-2 ECD/ligand complex showed that of the seven immunoglobulin-homology domains (Ig-domains) comprising the receptor ECD, Ig-domains 1-3 form the ligand binding site, while domains 4-7 are involved in homotypic receptor contacts, presumably regulating receptor dimerization and activation (Ruch et al., 2007; Stuttfeld and Ballmer-Hofer, 2009). In addition, high resolution information for Ig-domains D2-3 and D1-2 of VEGFR-2 and −3, respectively, bound to VEGF is available from our earlier work (Leppänen et al., 2013; Brozzo et al., 2012; Leppänen et al., 2010). These structures show the contribution of Ig-domains 1-3 of the receptors in ligand binding and how they interact with three variable loops present in all VEGF family ligands. In addition, a recent crystal structure of VEGFR-3 Ig-domains 4-5 indicated receptor-receptor interactions in D5 (Leppänen et al., 2013). Since the D5 interactions were a consequence of crystal packing, it has been left ambiguous whether these interactions represent the ligand induced homotypic contacts. To close this gap, we focused on the structural characterization of full-length VEGF receptor ECD complexes and finally succeeded in determining the structure of the full-length VEGFR-1 ECD in complex with VEGF-A at 4 Å resolution. The structure allows for the first time detailed visualization of ligand binding and ligand induced homotypic interactions within an entire VEGF receptor ECD/VEGF complex and opens new possibilities for future drug design.

## RESULTS

### Structure determination of the VEGFR-1 extracellular domain/VEGF-A complex

To obtain detailed structural information for future drug design we determined the organization of the full-length ECD of VEGFR-1 in complex with VEGF-A using X-ray crystallography. Human VEGFR-1 baculovirus constructs encompassing Ig-domains D1-7, D1-6, and D2-7 (Figure 1A) were expressed in insect cells, purified, and crystallized in complex with VEGF-A. Complex formation between ligand and receptor ECD proteins and monodispersity of the complexes was determined by size exclusion chromatography coupled to multi-angle light scattering (SEC-MALS) and small angle X-ray scattering (SAXS) (Supplemental Methods). Complex formation was already observed during protein purification as VEGF-A co-eluted from a SEC column together with the VEGFR-1 ECD (Figure S1A) and molecular weight determination by SEC-MALS clearly indicated 1:1 dimer formation of the full-length VEGFR-1 ECD with VEGF-A (Figure S1B). In addition, small angle X-ray scattering analysis in solution (SAXS) showed an increase in the radius of gyration (Rg) and of the maximal length of the complex (D_max_) in the presence of ligand as observed in the distance distribution function, confirming the presence of a stable ligand/receptor complex (Figure S1C).

**Figure 1.**
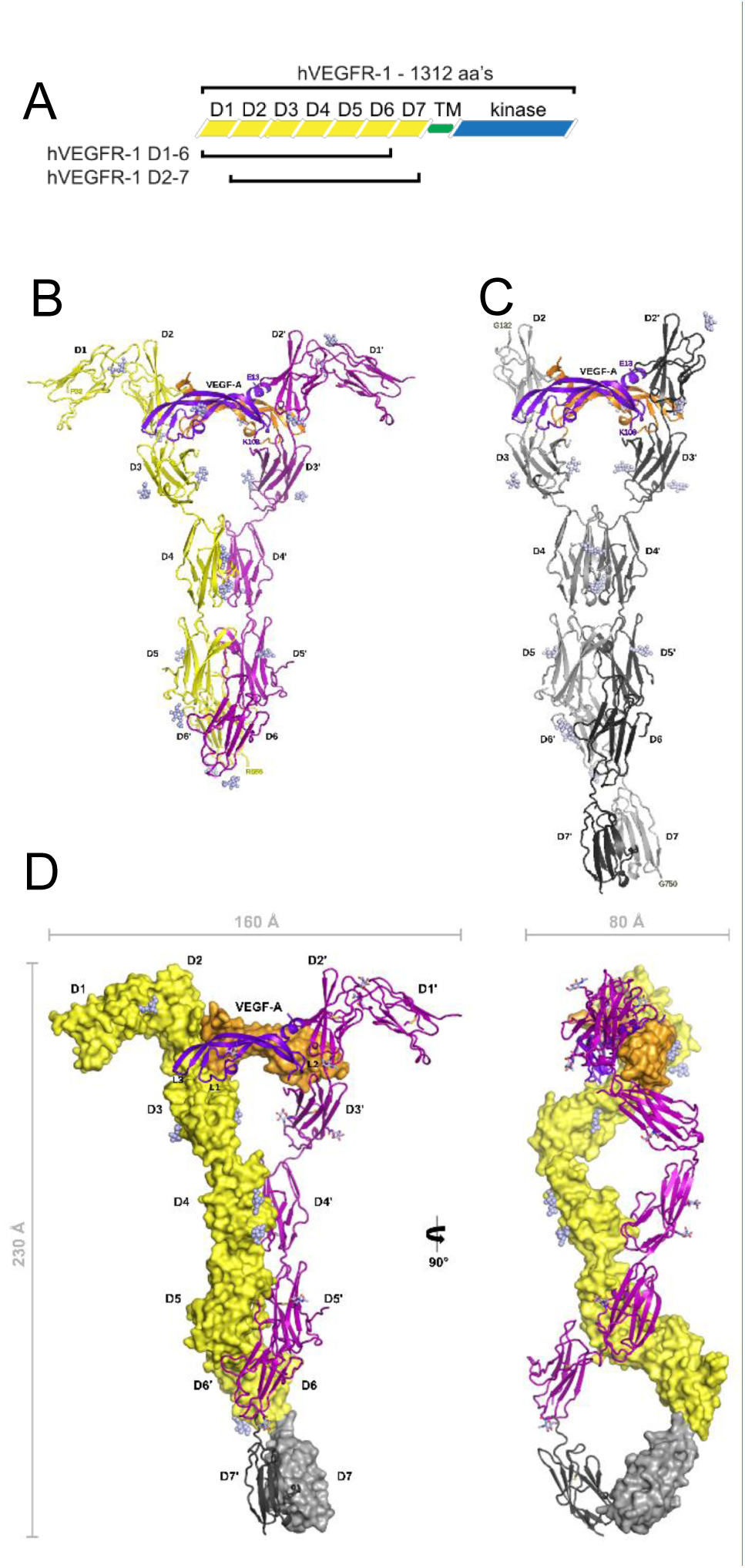
Structure of the VEGFR-1 extracellular domain in complex with VEGF-A. (A) Schematic overview of the domain organization of VEGFR-1. Crystallized constructs are indicated. Cartoon representation of the VEGFR-1 D1-6/VEGF-A complex (B) and of the VEGFR-1 D2-7/VEGF-A complex (C) structures. The chains of the VEGF-A homodimer are shown in purple and orange, the two chains of VEGFR-1 D1-6 in yellow and magenta, and the VEGFR-1 D2-7 chains in light and dark gray. N-linked glycans are depicted as spheres. (D) Composite model of the VEGFR-1 ECD/VEGF-A complex. One half of the complex is represented as surface with N-linked glycans depicted as spheres and the other half as cartoon with N-linked glycans depicted as sticks. The D7 homodimer shown in gray was positioned by superimposing VEGFR-1 D2-7/VEGF-A onto VEGFR-1 D1-6/VEGF-A. The dimensions of the complex and the axis of rotation relating the two views are also shown.

After various optimization rounds we obtained crystals of the D1-6 complex diffracting to 4 Å, which we found suitable for structure determination (Table 1). Crystals belonged to space group C2 with one complex per asymmetric unit and a solvent content of 70%. Enabled by a novel data collection strategy (Weinert et al., 2015), the structure of the VEGFR-1 D1-6/VEGF-A complex was solved by a combination of molecular replacement and single wavelength anomalous dispersion (MR-SAD) using iodide-soaked crystals (Figure 1B). The resulting electron density map revealed the D2 region and the ligand, positioned by molecular replacement, and contained continuous electron density for the missing Ig-domains (Figure S2). D3, D4, and D5 were placed by additional rounds of phased molecular replacement, and proper sequence assignment was performed by replacing the molecular replacement search models with the corresponding homology models generated by the SWISS-MODEL homology modeling server. D1 and D6, because of their less well-defined electron density and the lack of good homology modeling templates, were modeled using the Foldit structure prediction game (Cooper et al., 2010). The best D1 and D6 models were selected for further manual refinement based on their performance as search models in molecular replacement (Figure S3).

**Table 1.**
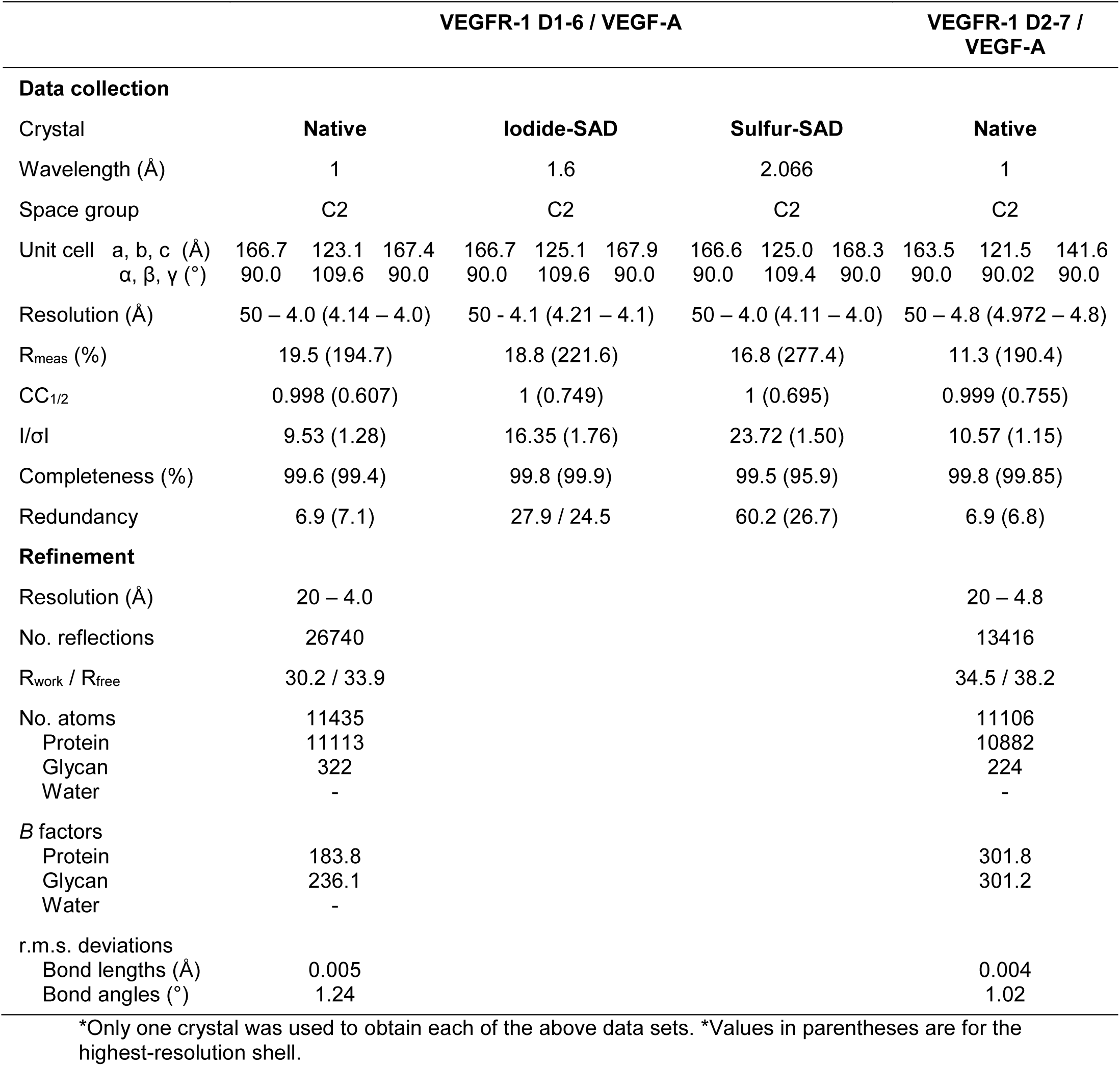
Data collection and refinement statistics

Anomalous scattering of Selenium-Methionine (Se-Met) is a powerful structure and sequence validation tool. In the absence of diffracting Se-Met derivative crystals we turned to sulfur-SAD to validate our final model of the VEGFR-1 D1-6/VEGF-A complex (Figure 2). With weakly diffracting crystals, careful data collection was crucial to obtain the accuracy necessary for resolution of the small anomalous signals. Despite the crystals diffracting only to 4 Å resolution, more than half of all anomalous scatterers, including every disulfide bridge in the Ig-domains and the VEGF-A ligand, as well as several methionines were assigned in the anomalous difference Fourier maps, verifying subunit placement as well as sequence assignment (Figure 2B). In addition, positive electron density present in the Fo-Fc maps for N-linked glycans around the asparagine residues predicted to be glycosylated further aided in exact sequence assignment.

**Figure 2.**
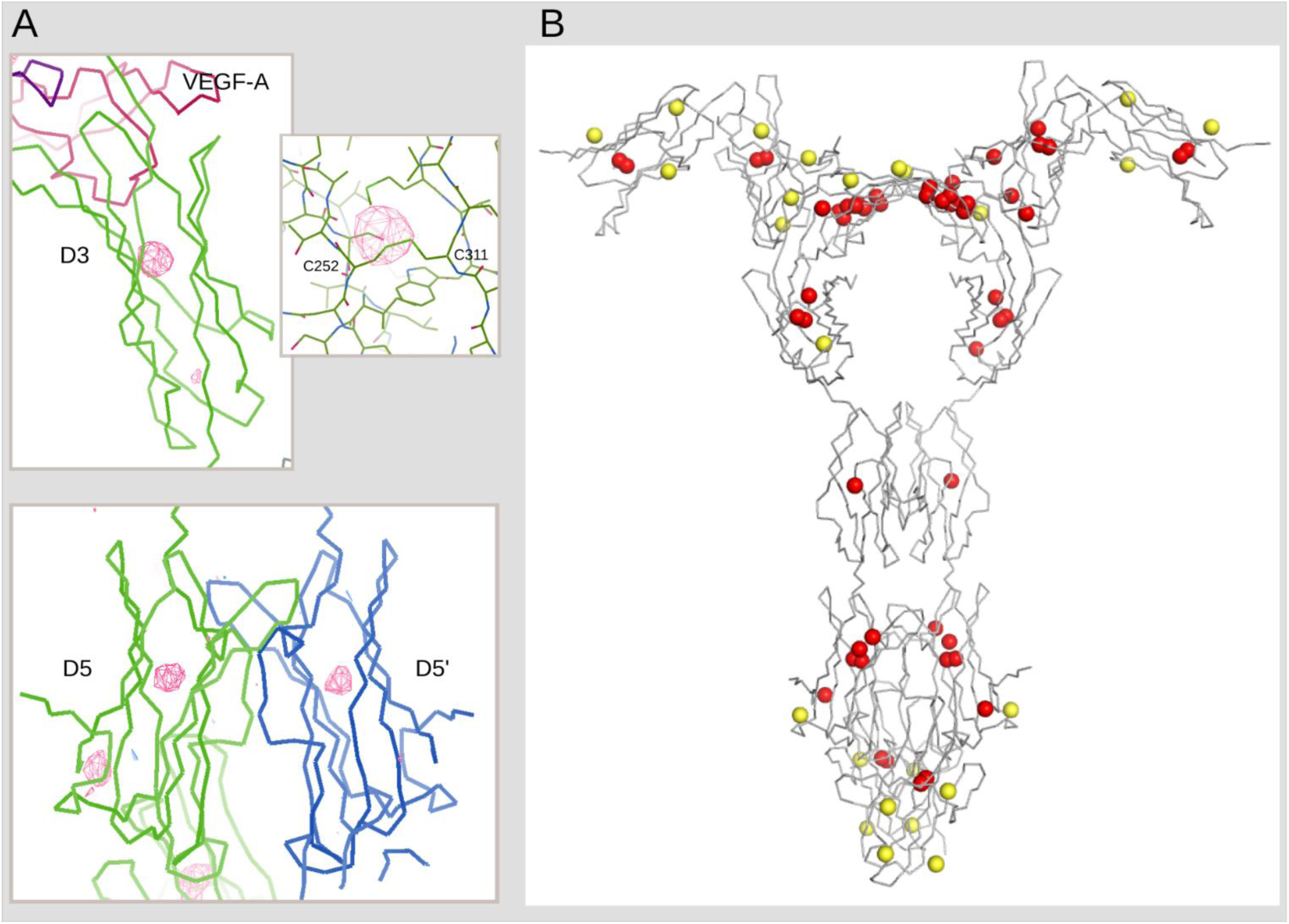
Native-SAD as validation tool. Ribbon representation of VEGFR-1 D3 (upper panel) and VEGFR-1 D5 dimer (lower panel) with sulfur peaks detected in anomalous difference Fourier maps. The map is contoured to 4σ. (B) Ribbon representation of the VEGFR-1 D1-6/VEGF-A complex structure with 86 sulfur atoms presented as spheres. 51 sulfur atoms (60%) colored in red were detected at 3σ in anomalous difference Fourier maps. The sulfur atoms colored in yellow belong mostly to methionines and were not detected.

To complete the molecular structure of the VEGFR-1 ECD, we also used crystals of the VEGFR-1 D2-7/VEGF-A complex diffracting to 4.8 Å. The D1-6 complex structure was used as a search model for molecular replacement of the D2-7/VEGF-A complex (Figure 1C). To create a composite model of the D1-7/VEGF-A complex we superimposed the two structures, D1-6/VEGF-A and D2-7/VEGF-A (RMSD of 1.76 Å for 1128 residues of the D2-6 region) and connected the D7 portion of the D2-7/VEGF-A model with the D1-6/VEGF-A based model (Figure 1D).

### Overall architecture of the VEGFR-1 ECD/VEGF-A complex

The structures of the D1-6 and the D2-7 VEGF-A complexes both show 2:2 stoichiometry and contain two sets of 1:1 complexes in the asymmetric unit related by 2-fold non-crystallographic symmetry. In the composite model of the VEGFR-1 ECD/VEGF-A complex (Figure 1D), two receptor chains are bridged with the dimeric VEGF-A ligand in the D2-3 region and related by two-fold symmetry throughout the long axis of the complex. The complex has overall dimensions of ~160 Å × ~230 Å × ~80 Å and is characterized by the existence of three cavities: the upper one formed between the VEGF-A dimer, D3 and D4 domains of two receptor chains (dimensions ~65 Å × ~50 Å); the middle one between the D4 and D5 domains (~30 Å × ~15 Å); and the lower one between the D5, D6 and D7 domains (~55 Å × ~50 Å). VEGF-A binding to D2 and D3 induces a twisted arrangement between these two domains, with a bending angle of ~135°. This is followed by intertwining of receptor chains in D4-5 bringing two protomers in close proximity to each other. The twist angle between D4 and D5 within the same chain is 60°. Additional homotypic contacts are observed between D7 domains. In our model the D7 dimer deviates from the D1-6/VEGF-A 2-fold axis, which is a consequence of crystal packing of the D2-7/VEGF-A complex. This finding suggests flexibility in the D6-7 region due to a longer linker between these two domains.

To confirm the overall structure of the VEGFR-1 ECD/VEGF-A complex we additionally visualized the complex by negative stain electron microscopy (Figure 3A). The calculated class averages showed excellent agreement with projections of the crystal structure (Figure 3B). All class averages show dimeric VEGFR-1 ECD chains bridged by VEGF-A. Interestingly, a slight bend is observed in the membrane-proximal region confirming the kinked D6-D7 region seen in the D2-7/VEGF-A crystal structure. As there is a clear preference for the complex to adsorb to the grid showing the front view, only one class was generated of the side view.

**Figure 3.**
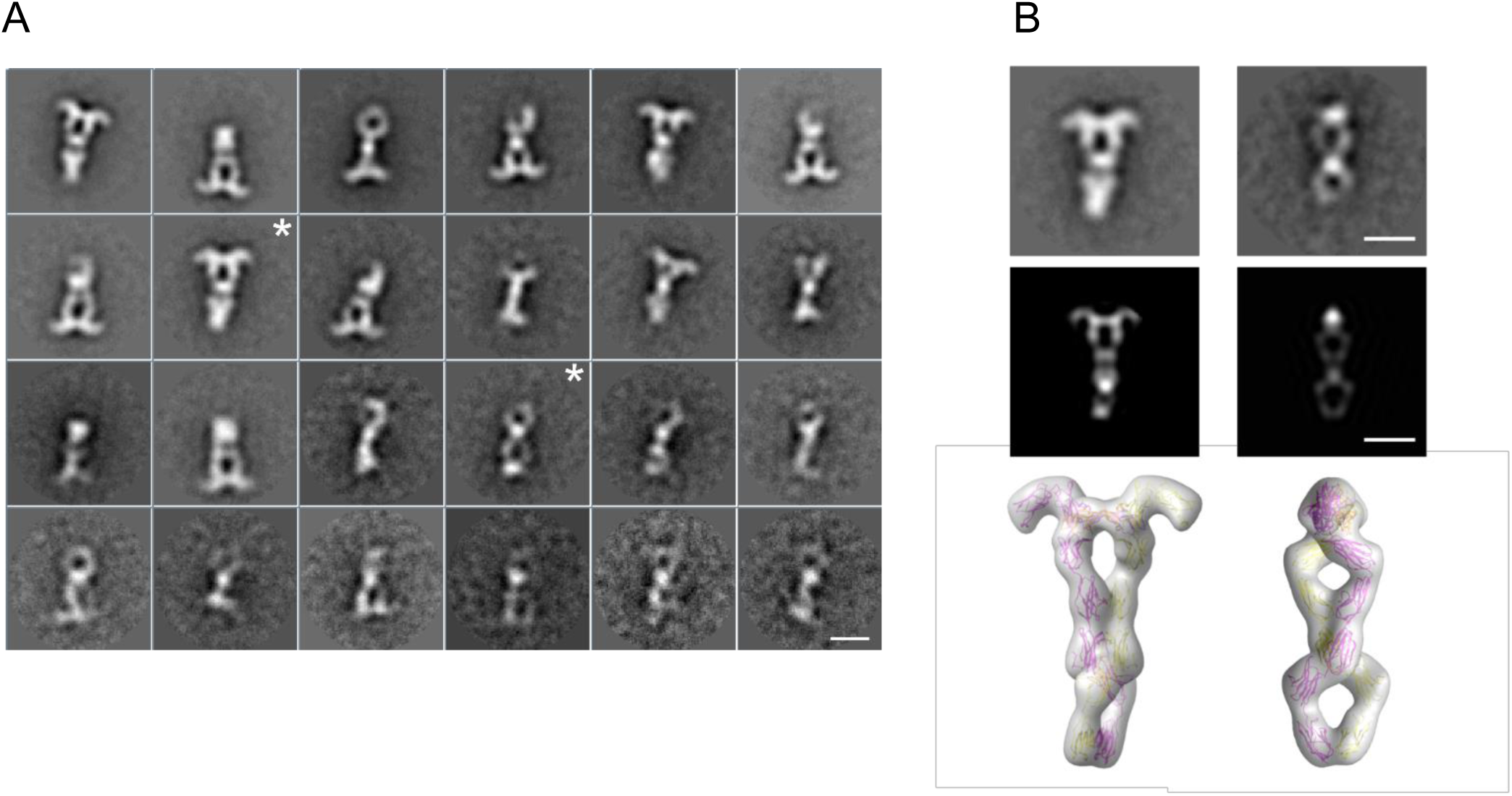
Negative-staining EM analysis of the VEGFR-1 D1-7/VEGF-A complex. (A)A gallery of 24 2D class averages from image classification is shown. (B) Comparison of the VEGFR-1 D1-7/VEGF-A EM class averages with the composite model shown in Fig. 1. Representative class averages (labeled with a star in A) are compared with selected 2D projections and with the corresponding 3D-volume models of the VEGFR-1 D1-7/VEGF-A complex. The volumes were filtered at 25 Å resolution. The class average in the right panel of b was rotated by ~180° relative to its orientation in a (Scale bars: 10 nm).

### Individual Ig-domains are all glycosylated and apart from D6 belong to the I-set of the Ig-domain superfamily

This is the largest structure determined so far in the VEGF receptor family and allows us to describe the structure and arrangement of the individual Ig-domains of VEGFR-With the exception of D6, all Ig-domains of VEGFR-1 belong to the I-set of the Ig-domain superfamily. D1 (residues 32-130) was modeled by Foldit players and subsequently rebuilt during refinement. Although the experimental electron density map suggested large disordered regions, the 2Fo-Fc maps allowed us to build all residues except the D-E loop (78-84), which was disordered. Additional electron density around N100 confirmed the presence of an N-linked glycan, and the position of a disulfide bridge between C53 and C107 was validated by sulfur-SAD experiments. The structure of D2 (residues 132-225) was solved before (Wiesmann et al., 1997) and our structure additionally shows N-glycosylation of N164 and N196. D1 is bent away by ~ 80° from D2 and has no contact with the ligand. The bent conformations of the D1-2 modules of VEGFR-1 and VEGFR-3 (Leppänen et al., 2013) are very similar and can be aligned with an RMSD of 1.66 Å for 161 residues (Figure S4). D3 (residues 226-330) is an elongated Ig-domain with N-glycosylation of N251 and N323. D4 (residues 333-426) is a smaller Ig-domain not disulfide bridged between the β-sheets and lacking β-strand 4. D4 is glycosylated at N402 and N417. D5 (residues 428-554) is the largest Ig-domain in VEGF receptors due to a long C-D loop that connects two β-sheets. Residues 472-486 within this region were disordered and therefore omitted from the final model. In D5 we observed N-glycosylation of N547 and verified the positions of the disulfide bridge between C454 and C535 and of all methionines by S-SAD (Figure 2). With our structure we obtained, for the first time, structural information on D6 of VEGF receptors, which belongs to the C2-set of Ig-domains. D6 contains strands of intermediate length in both β-sheets, a disulfide bridge between C577 and C636, and an N-linked glycan at N625. D7 (residues 661-750) was modeled using D7 of VEGFR-2 (PDB ID: 3KVQ) as a template and, due to the low resolution of the VEGFR-1 D2-7 complex data, we do not further discuss its structural details. However, D7 of the two protomers form a dimeric arrangement as described in the crystal structure of isolated VEGFR-2 D7 (Yang et al., 2010).

### VEGFR-1 D3 contributes to ligand binding and determines ligand specificity

Ligand binding to VEGFR-1 takes place in Ig-domains 2 and 3. With our structure we can now describe the complete ligand binding site of VEGFR-1, while earlier published complex structures contained only D2 (Christinger et al., 2004; Iyer et al., 2010; Wiesmann et al., 1997). Each receptor chain interacts with both VEGF-A protomers (Figure 4A). The surface buried on one receptor chain by the bound VEGF-A dimer is ~1500 Å^2^, with D2 and D3 contributing similar interface areas (~800 Å^2^ for D2 versus ~700 Å^2^ for D3). VEGF-A residues interacting with D2 are part of the N-terminal helix α1 of the ligand protomer A (M17, F18, Y21, Q22 and Y25); and of the strands β2 (I46, K48), β4 (Q79, M81, I83) and β5 (Q89, I91) of the protomer B (Figure 4B). While the interface between ligand and D2 is largely hydrophobic, the contacts with D3 are hydrophilic comprising a number of ionic interactions. There is an overall positive charge of the D3 region that comes in contact with the ligand. Several charged residues from all three loops (L1, L2 and L3) of VEGF-A interact with D3. The most prominent conserved residue is E64 in L2 that engages 5 charged residues located in D3 of the receptor (N259, R261, R280, Q284 and N290) in hydrogen bond and salt bridge type interactions. Another residue from L2, D63, forms salt bridges with R224 located in the D2-3 linker (Figure 4C). In addition, E44 in L1 and K84 in L3 form hydrogen bonds with Q263 of D3. Loops 1 and 3 also carry hydrophobic residues such as P40, I43 and P85 that form complementary surface with D3 of VEGFR-1. I43 protrudes into the hydrophobic pocket on the D3 surface formed by V262, M264, V278 and F292.

**Figure 4.**
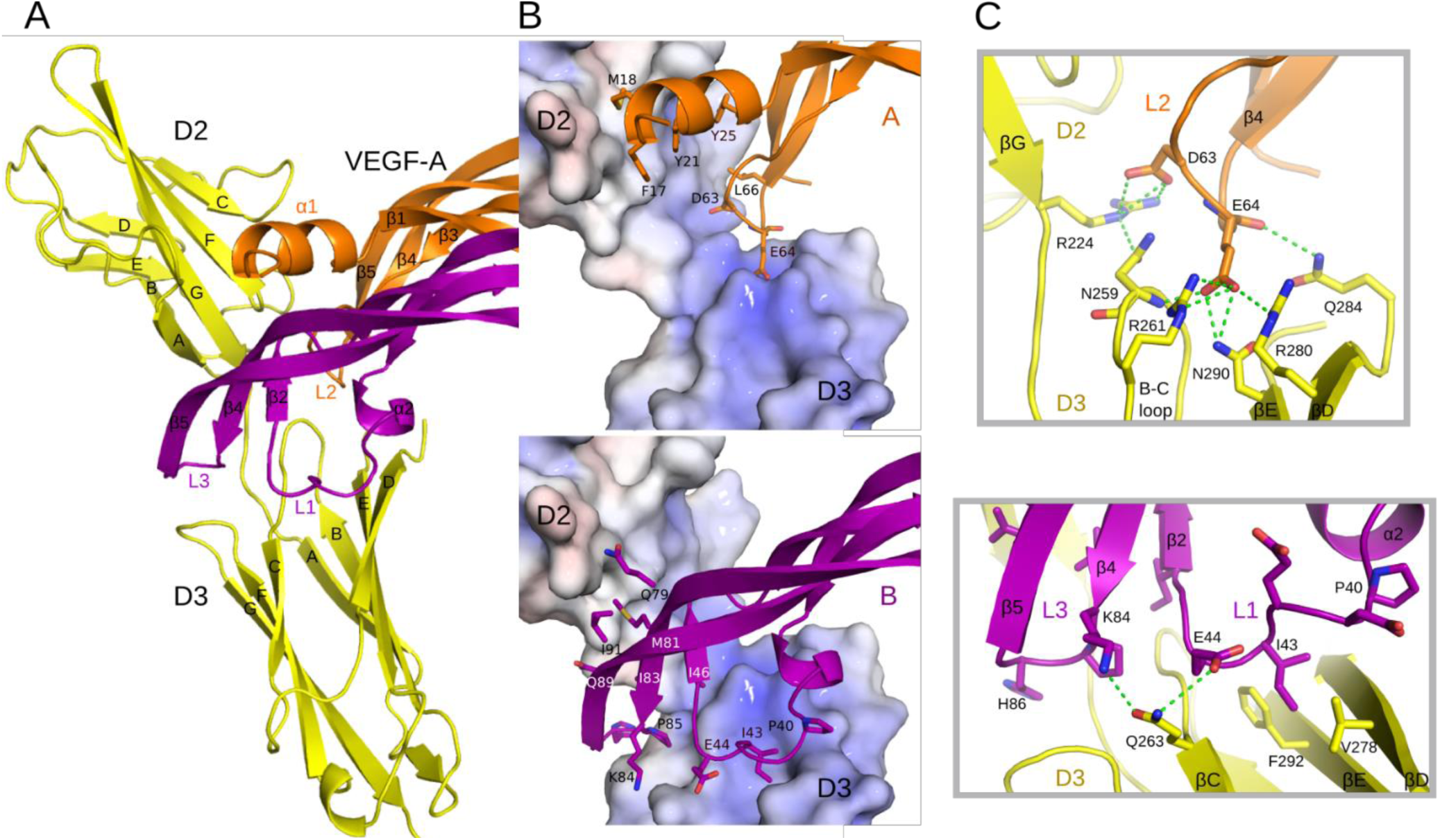
Binding interface between VEGF-A and VEGFR-1 D2-3. Cartoon representation of D2 and D3 of one receptor chain colored in yellow and VEGF-A chains in orange (chain A) and purple (chain B). Secondary structure elements are labeled. Three loops in VEGF-A are designated L1, L2 and L3. (B) VEGFR-1 interaction with the VEGF-A monomer A including N-terminal helix α1 and loop 2 (upper panel) and with the VEGF-A monomer B including loops 1 and 3 (lower panel). The key residues of the ligand are highlighted as sticks and labeled. VEGFR-1 charge distribution at the interaction surface is presented as a surface potential model (calculated with the APBS module in PyMol). (C) Interactions of charged residues in VEGF-A L2 (upper panel) and L1 and L3 (lower panel) with VEGFR-1 D3. Hydrogen bonds and salt bridges are shown as green dashed lines.

Structural comparison of receptor binding epitopes for the VEGFR-1 ligands VEGF-A, PlGF and VEGF-B shows that all three ligands interact similarly with D2 of VEGFR-1 (Figure S5). The interaction between the loop L2 of the three ligands with D3 is also very similar, since D63 of VEGF-A is strictly conserved and E64 is conserved in PlGF and substituted by aspartate in VEGF-B. Differences are observed in the way how the three ligands interact through the loops L1 and L3 with D3 of VEGFR-1. While the sequence and structure conservation in these regions is high between VEGF-A and PlGF, this is not the case for VEGF-B. Superposition of the VEGFR-1 D2/VEGF-B complex onto our structure enables visualization of the possible structural arrangement of the VEGF-B loops L1 and L3 in respect to receptor domain D3. It appears that these loops are moved away from D3 and not involved in any interactions. These findings agree with a recent study (Anisimov et al., 2013) that showed that VEGF-B does not require D3 interactions for high-affinity binding and does not promote signaling downstream of VEGFR-1. PlGF and VEGF-A, on the other hand, are angiogenic ligands and the L1 interaction with D3 is important for receptor activation.

### VEGFR-1 D4-5 form homotypic interactions in the presence of ligand

The structure of the VEGFR-1 D1-6/VEGF-A complex reveals homotypic interactions in D4 and D5, covering altogether a solvent-accessible area of ~1000 Å^2^ per chain (Figure 5A and B). The interface between the Ig-domains 4 is small (~165 Å^2^ per chain) and includes R351 located in the A-B loop of one D4 that shares a hydrogen bond with S380 located in the C-E loop of the other D4 (Figure 5C). The interactions between the residues in the E-F loop proposed earlier based on a KIT structure (Yuzawa et al., 2007) are not present in our structure.

**Figure 5.**
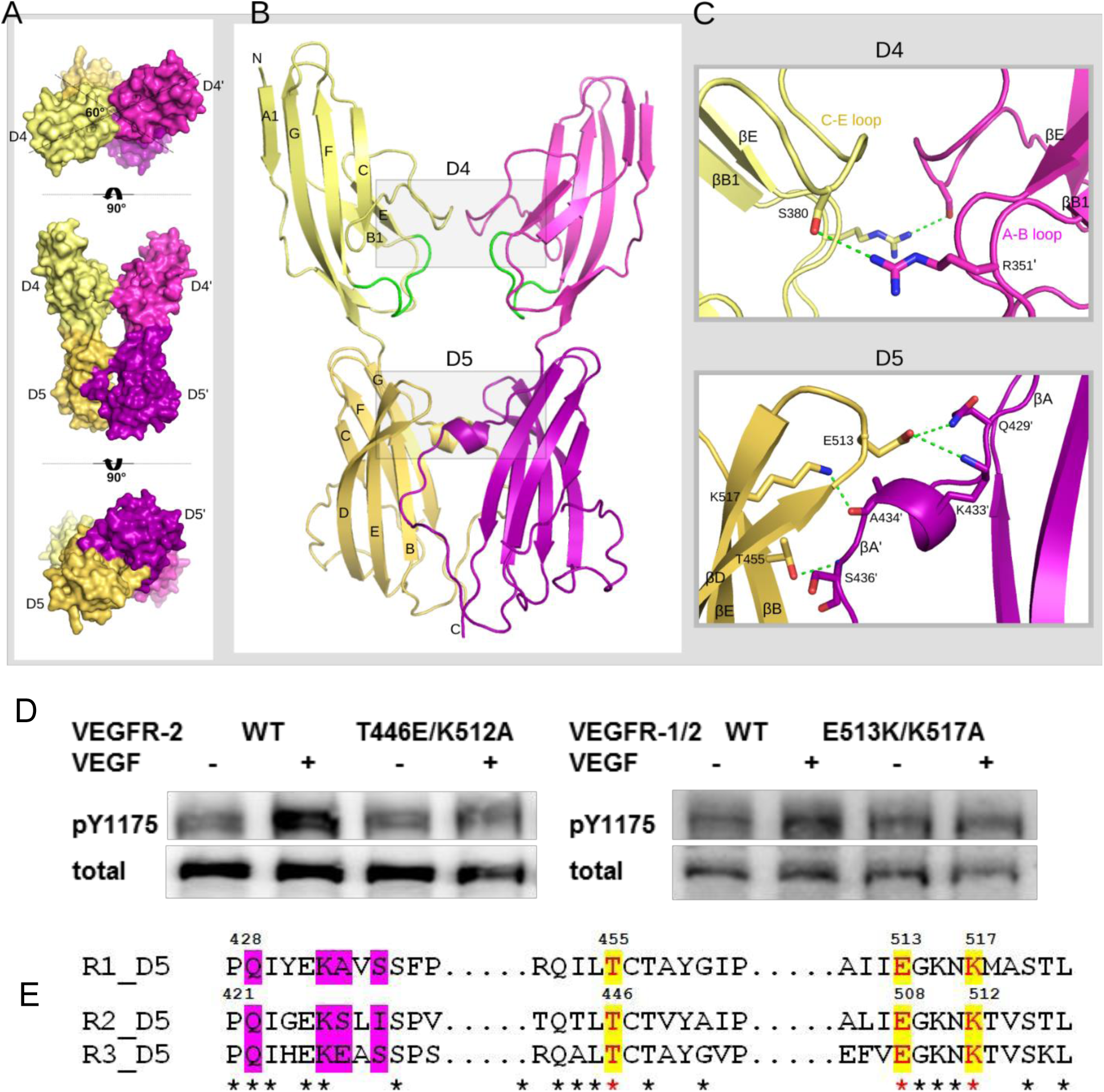
Homotypic interactions between Ig-domains 4 and 5 of VEGFR-1. (A) Surface representation of the D4-5 dimer. D4 on two chains are colored in pale-yellow and light-magenta and D5 is shown in yellow-orange and purple. The rotation axis relating the views is shown. The twist angle between D4 and D5 within the same chain is 60°. (B) Cartoon representation of the D4-5 dimer with labeled β-strands. The E-F loop in D4 is colored in green illustrating that residues that are part of this loop in VEGFR-1 do not interact. (C) Zoom-in view into the interface of D4 (top) and D5 (bottom). Interacting residues are shown as sticks and hydrogen bonds and salt bridges as green dashed lines. (D) Ligand-mediated receptor activation of wt and VEGFR D5 mutant receptors. Left panel shows Western blots for wt and mutant T446E/K512A VEGFR-2, right panel for wt and mutant E513K/K517A chimeric VEGFR-1/2 receptor consisting of the ECD of VEGFR-1 and the kinase domain of VEGFR-2. Activities were determined with phospho-specific antibodies as described in Supplemental Methods. (E) Sequence alignment of the subdomain of D5 involved in homotypic contacts in VEGFR-1, −2 and −3.

The homotypic contacts in D5 are, like in VEGFR-3, centered on the fully conserved residues T455 (β-strand B) and K517 (β-strand E) that form hydrogen bonds with the backbone atoms of S436 and A434 of the other chain (Figure 5C). Furthermore, a glutamate residue in the D-E loop (E513) conserved in all VEGFRs interacts *via* hydrogen bond with Q429 and *via* salt bridge with K433 of the other chain (Figure 5E). While the fully conserved residues are part of the rigid β-sheet, their interacting partners are not conserved and belong to the flexible strands A and A’ and the helical protrusion between them.

### Homotypic interactions contribute to ligand affinity and full receptor activity

The purified receptor ECD was monomeric indicating that the homotypic interactions in D4-7 are weak and apparently depend on ligand-mediated receptor dimerization (Figure S1). We used isothermal titration calorimetry (ITC) to assess the contribution of the membrane-proximal Ig-domains 4-7 to the binding affinity of the full-length ECD of VEGFR-1 for VEGF-A. Similar to VEGFR-3 (Leppänen et al., 2013), D1-7 of VEGFR-1 had significantly higher affinity for the ligand than D1-3 (Figure 6A). The binding affinity of VEGFR-1 D1-7 for VEGF-A was in the low nanomolar range (~ 2 nM), while the affinity of VEGF-A for VEGFR-1 D1-3 was 20 times lower i.e. ~ 50 nM. Binding of VEGF-A to D1-3 was entropically favored, whilst binding to D1-7 was enthalpically driven (Figure 6B). The presence of Ig-domains 4-7 thus increased binding affinity for VEGF-A, presumably due to stabilization through homotypic receptor-receptor contacts in D4-7.

**Figure 6.**
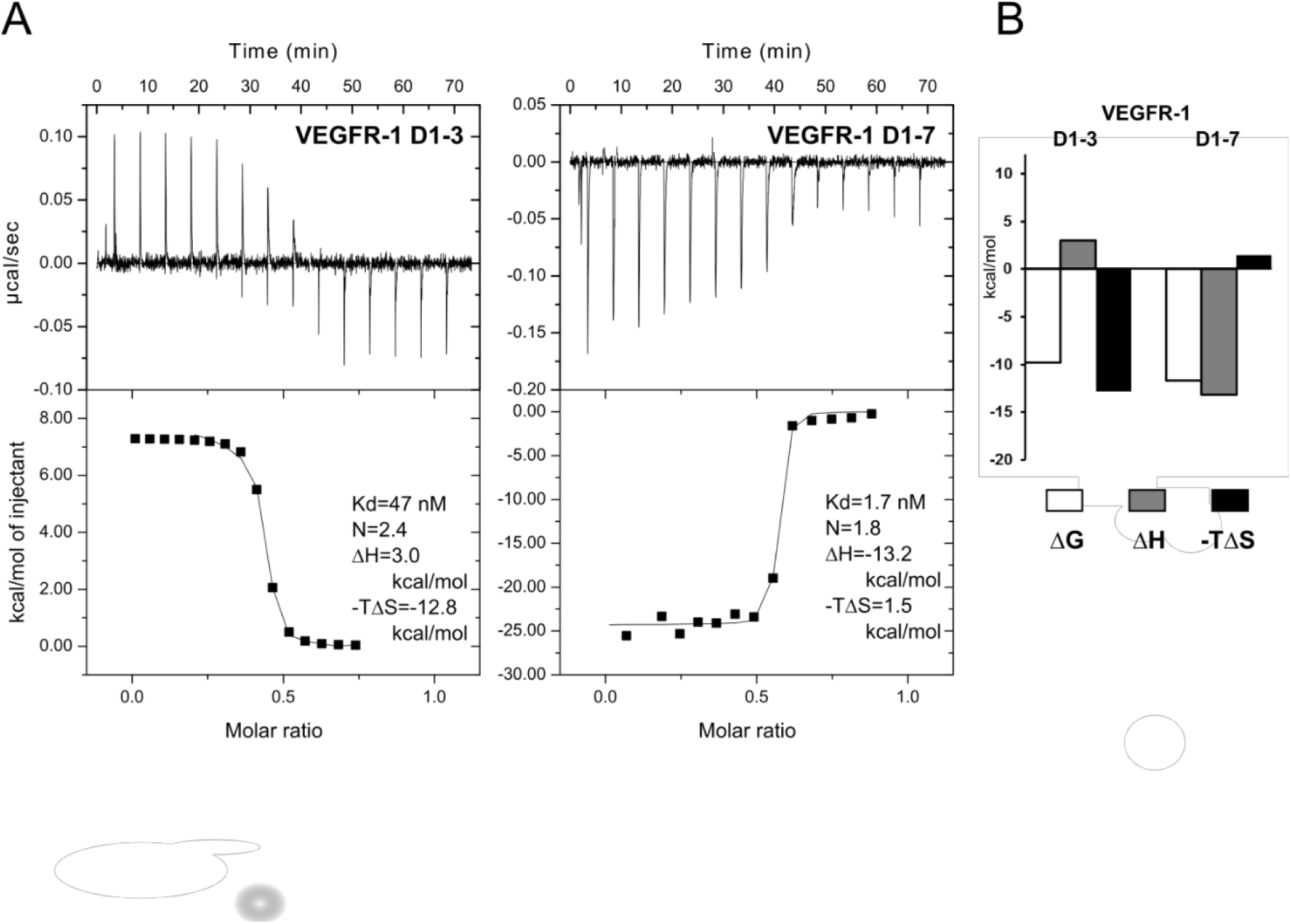
Quantification of VEGFR-1/VEGF-A interaction by Isothermal TitrationCalorimetry (ITC). (A) Raw titration data and integrated and concentration normalized isothermograms of VEGF-A binding to the VEGFR-1 D1-3 (left panels) and VEGFR-1 D1-7 (right panels). Solid lines in the isothermograms represent the best fit according to the “One Set of Sites” model. (B) Thermodynamic signature of complex formation-enthalpy and entropy contribution to the Gibbs free energy of binding-differs between the two constructs of the receptor.

Based on the structural data described here and in our earlier work (Leppänen et al., 2013; Brozzo et al., 2012; Leppänen et al., 2010; Ruch et al., 2007; Kisko et al., 2011) we predicted that the contacts in D5 are essential for receptor function. We therefore mutated the residues involved in the D5 interaction and determined receptor activity in transfected NIH 3T3 cells. The activity of VEGFR-1 was too low to obtain high quality functional data and we therefore mutated the corresponding residues in D5 of VEGFR-2 based on a homology model of this domain (for details of the mutants used see Figure 5E). In the mutant T446E/K512A residue T446 was replaced by E and K512 by A, thus preventing hydrogen bond and salt bridge formation in the dimeric receptor complex. This mutant showed a decrease in ligand-induced VEGFR-2 phosphorylation by 60% reflecting impaired receptor activation (Figure 5D left panel). To determine the functionality of the VEGFR-1 D5 interface we also created a chimeric VEGFR (VEGFR-1/2) consisting of the VEGFR-1 ECD and the intracellular domain of VEGFR-2. Mutant E513K/K517A showed a reduction in receptor kinase activity by 50% (Figure 5D right panel). Taken together, these data show that the contacts in D5 revealed by our VEGFR-1 ECD/VEGF-A structure are essential for receptor activation.

## DISCUSSION

We present here the first full-length VEGF receptor ECD structure in complex with one of its ligands revealing all structural details of VEGF-A binding to VEGFR-1. Most importantly, our structure shows the structural details of the homotypic contacts in D4-7 in the context of the full-length receptor ECD. Our functional analysis of these interactions proves their relevance for receptor activation. We propose that exact positioning of ligand-bound receptor protomers in active dimers, which is essential for kinase activation, is mediated by the concerted interplay of ligand-receptor interactions in D2-3 and receptor-receptor interactions in D4, D5 and D7.

### Specificity of ligand binding to VEGFRs

Although earlier studies did not report increased affinity towards VEGF-A when including Ig-domain 3 in VEGFR-1 constructs (Wiesmann et al., 1997), our structure shows significant D3 interactions with all three connecting loops of VEGF-A. We propose that the role of D3/VEGF interactions is to trigger receptor ECD intertwining and to induce homotypic interactions in D4-5 and D7. In addition, we identify D3/VEGF interaction as a determinant for ligand specificity and as a sensor for modification of homotypic interactions in D4-7 depending on the strength of the induced twist between D2-3. This might ultimately lead to a different functional output induced by distinct VEGF ligands. With the structure of the VEGFR-1 ECD/VEGF-A complex and the comparison of receptor binding epitopes in VEGF-A, PlGF and VEGF-B we provide a structural basis for understanding the different signaling outputs generated by these three VEGFR-1 ligands. Our data thus provide structural evidence for the role of the VEGFR-1 D3/ligand interface in receptor signaling (Figure S5).

The next question that arises is why do VEGF-C,-D and-E not bind to VEGFR-1? Sequence conservation between different VEGF family members varies in the three binding loops. While the residues in the loop L2 are highly conserved within the VEGF family, there are significant sequence and structural differences in the N-terminal helix α1 and in the loops L1 and L3, which presumably define receptor specificity of the ligands. The interaction between residues E44 in L1 and K84 in L3 of VEGF-A with D3 of VEGFR-1 are not conserved in VEGF-C, D, and E (Figure S6), and may thus determine receptor specificity.

Structural comparison of VEGF-A binding to VEGFR-1, as described here, with binding to VEGFR-2 (PDB ID: 3V2A) shows similar type of interactions. The ligands in the two structures have very similar structures and can be aligned with an RMSD of 0.86 Å for 190 aligned residues. The difference exists in the orientation of D3 towards D2 where VEGF-A binding to VEGFR-1 induces a stronger twist by 11° between D2 and D3 (Figure S4). With the exception of E64 in VEGFR-1, which engages more residues in salt bridge and hydrogen-bond type interactions, loop L2 interacts similarly with D3 of VEGFR-1 or VEGFR-2.

To fully understand the differences in the signaling output by VEGFRs upon binding to a particular VEGF ligand, one needs to consider also the role of co-receptors. Distinct ligands promote ternary complex formation, which has an impact on receptor trafficking and signaling (Ballmer-Hofer et al., 2011).

### Role of homotypic receptor contacts in VEGFR activation

Our structural analysis describes for the first time the details of homotypic contacts in the membrane proximal domain of type V RTKs in the context of a full-length ligand-bound receptor ECD. In addition, we show that mutation of the interacting interface attenuates receptor activation. This is in agreement with earlier findings showing that homotypic contacts in D4-7 are functionally relevant for VEGFR-3 (Leppänen et al., 2013) and VEGFR-2 (Yang et al., 2010) and documents the strong similarity with type III RTKs such as PDGFR (Yang et al., 2008) and Kit (Yuzawa et al., 2007; Reshetnyak et al., 2015; Yang et al., 2010). The described ectodomain contacts in the membrane proximal domain are apparently essential for the tight control of type III and V receptors. Mutation of the interface involved in these homotypic contacts renders e.g. KIT oncogenic (Reshetnyak et al., 2015) and promotes ligand-independent kinase activation. We also showed earlier that antibody like molecules such as DARPins (Stumpp et al., 2008; Binz et al., 2004) binding D4 or D7 of VEGFR-2, block receptor signaling (Hyde et al., 2012).

Receptor-receptor interactions in D4 revealed in this work are in agreement with earlier studies on VEGFR-1 (Barleon et al., 1997) and VEGFR-2 (Shinkai et al., 1998) that pointed at the importance of this domain in ligand-induced receptor dimerization. Similar interactions were, on the other hand, not observed in the VEGFR-3 D4-5 structure (Leppänen et al., 2013). Since the D4 interactions in VEGFR-1 occur through the C-E loop, which is longer in this receptor compared to the other homologs, they might be relevant for VEGFR-1 functionality. Interestingly, in our structure we did not observe the previously proposed D4 interactions between residues in the E-F loop (Yuzawa et al., 2007). However, careful analysis of the D4 residues facing each other reveals that R382 in the C-E loop and D394 in the E-F loop could, by simple modification of their chi angles, form a salt bridge. This putative D4 interface would be very similar to the interactions in D4 of KIT (Yuzawa et al., 2007) and D7 of VEGFR-2 (Yang et al., 2010). In the current structure R382 forms an internal salt bridge with D399. Whether the side chains of R382 and D394 may adopt multiple conformations cannot be determined at the resolution of our structure (Figure S7).

Homotypic interactions in D5 have interesting structural characteristics with one interacting side composed of a rigid β-sheet (β-strands B, D and E) and the other side being more flexible (strands A and A’) (
Figure 5C
). Curiously, the flexible strands A and A’ are directly linked to D4. This structural feature may suggest the order of homotypic contact formation following ligand binding, *i.e.* formation of interactions in D4 will be transmitted downstream to D5. It may also explain the possibility of heterodimer formation (VEGFR-1/VEGFR-2 and VEGFR-2/VEGFR-3) by having one interacting face rigid and fully conserved, and the other more flexible allowing for differential bond formation.

Finally, the D7 interactions initially observed in VEGFR-2 (Yang et al., 2010) and now confirmed for VEGFR-1 represent one more check-point in the formation of an active receptor dimer. This additional restriction of mobility towards the transmembrane domain is probably necessary for more stringent control of VEGFR activation compared to the type III RTK family members.

Taken together, the availability of the structural details of the full-length VEGFR-1 ECD in complex with VEGF-A presented here allow a detailed mechanistic interpretation of the role of individual Ig-domains in receptor dimerization and activation. Most drugs targeting VEGF receptors to interfere with aberrant vascularization available today are specific for VEGFR-2. The role of VEGFR-1 in angiogenesis, and particularly in tumor angiogenesis, has recently attracted high attention as it seems to be particularly relevant for *de novo* vascularization as demonstrated for instance by Massena et al. (Massena et al., 2015). Additional relevance of VEGFR-1 in pathological angiogenesis has been observed in cancer models (Van de Veire et al., 2010; Schomber et al., 2007). The discovery of the mechanism of receptor activity modulation by domains 4-7 described here opens new possibilities for developing novel, highly specific VEGF receptor antagonists for future medical applications.

## EXPERIMENTAL PROCEDURES

### Production and purification of recombinant proteins

Human VEGF-A_121_ (here denoted as VEGF-A) was produced in *Pichia pastoris* as described before (Scheidegger et al., 1999). Human VEGFR-1 D1-3 (residues 1-339), D1-6 (residues 1-660), D2-7 (residues 132-750) and D1-7 (residues 1-750) were expressed as secreted proteins in Sf21 insect cells, as described previously (Brozzo et al., 2012). The proteins were purified from culture supernatant by immobilized metal affinity chromatography (IMAC) followed by size exclusion chromatography (SEC) on Superdex 200 HR 16/600 (GE Healthcare) equilibrated in 25 mM HEPES pH 7.5 and 500 mM NaCl. For the formation of VEGFR-1 ECD/VEGF-A complexes, an equimolar amount of receptor and ligand were mixed and the complexes were purified by SEC.

### Size exclusion chromatography coupled to multi-angle light scattering (SEC-MALS)

Experimental details for SEC-MALS are given in Supplemental Experimental Procedures.

### Isothermal titration calorimetry (ITC)

Experimental details for ITC are given in Supplemental Experimental Procedures.

### Cell transfection and determination of receptor activity

NIH 3T3 cells grown in 6 cm plates in DMEM supplemented with 10% calf serum were transfected with VEGFR-2 expression constructs and processed as described in Supplemental Experimental Procedures.

### Negative stain single particle electron microscopy (EM)

Experimental details for negative stain EM are given in Supplemental Experimental Procedures.

### Comparison of the crystal and EM structure of the VEGFR-1 D1-7/VEGF-A complex

The composite model of the VEGFR-1 D1-7/VEGF-A complex was converted into density volume filtered to 25 Å resolution using the Bsoft software package (Heymann and Belnap, 2007). The volume was then used to calculate projections at an angular interval of 15° with the SPIDER image processing suite (Frank et al., 1996). The projections with the most similar features to the experimental 2D averages were selected. Density volumes and ribbon diagrams were displayed and oriented with the UCSF Chimera package (Pettersen et al., 2004).

### Small angle X-ray solution scattering (SAXS)

SAXS data acquisition was performed at the X12SA-beamline (cSAXS) at the Swiss Light Source at the Paul Scherrer Institute in Villigen, Switzerland, experimental details are given in Supplemental Experimental Procedures

### Crystallization and structure determination

*Crystallization.* The VEGFR-1 D1-6/VEGF-A and the VEGFR-1 D2-7/VEGF-A complexes used for crystallization were partially deglycosylated with PNG-ase F and purified by SEC. Both complexes were crystallized in sitting drops at 20°C. The best diffracting crystals of the VEGFR-1 D1-6/VEGF-A complex were obtained from a reservoir solution containing 200 mM CaCl_2_, 100 mM MES pH 6.5 and 10-14% PEG 3000 (w/v) and of the VEGFR-1 D2-7/VEGF-A complex from a reservoir solution containing 200 mM NaCl, 100 mM Na/K-phosphate pH 6.2 and 10% PEG 8000 (w/v). The crystals of both complexes belonged to space group C2 with one complex per asymmetric unit, a solvent content of 70% and the following cell parameters: VEGFR-D1-6/VEGF-A a=166.7, b=123.1, c=167.4 Å, β=109.6°; VEGFR-1 D2-7/VEGF-A a=163.5, b=121.5, c=141.6 Å, β=90.02°. Iodide derivatives of the VEGFR-1 D1-6/VEGF-A complex were prepared by soaking the crystals in a reservoir solution containing 100 mM CaCl_2_ for 3 hrs. For cryoprotection, the crystals were transferred stepwise into mother liquor containing increasing concentrations of glycerol to a final concentration of 20%, flash-frozen and stored in liquid nitrogen.

*Structure determination.* All datasets were collected on the X06DA beamline at the Swiss Light Source (SLS) at the Paul Scherrer Institute in Villigen, Switzerland on a PILATUS 2M detector (Dectris). A complete dataset to 4 Å resolution was collected from a single native crystal of the VEGFR-1 D1-6/VEGF-A complex. Highly redundant anomalous data were obtained from a single large crystal derivatized with iodide that was measured on multiple positions and orientations using the high-precision multi-axis goniometer PRIGo (Waltersperger et al., 2015).

Data were indexed, integrated and scaled with XDS (Kabsch, 1993) and further processed with CCP4 programs (Collaborative Computational Project, 1994). The structure of the VEGFR-1 D1-6/VEGF-A complex was determined by a combination of molecular replacement and single-wavelength anomalous dispersion (MR-SAD), as implemented in the program suite PHASER (McCoy et al., 2007). The partial molecular replacement solution obtained using the structure of the VEGFR-1 D2/VEGF-A complex (PDB ID: 1QTY) as a search model allowed for the identification of heavy atom sites and substructure completion. Initial phases were improved by solvent flattening and extended to 4 Å by two-fold NCS averaging with the program DM (Cowtan et al., 2012). The missing Ig-homology domains were positioned by phased molecular replacement in MOLREP (Vagin and Teplyakov, 2010) using the following structures as search models in the following order: for D4 VEGFR-3 D4 (PDB ID: 4BSJ), for D5 VEGFR-3 D5 (PDB ID: 4BSJ) and for D3 VEGFR-2 D3 (PDB ID: 2X1X). To construct a model with the full VEGFR-1 sequence we created homology models for the Ig-domains D3-D5 using the corresponding MR search models as templates in SWISS-MODEL (Biasini et al., 2014). The homology models were superimposed onto the domains positioned by molecular replacement and single Ig-domains were connected generating the VEGFR-1 D2-D5/VEGF-A model.

We first attempted to fit D1 and D6 using homology models as well, but the resulting models had a poor fit without clear difference density to guide model building. To overcome this, D1 and D6 were submitted to the Foldit structure prediction game (Cooper et al., 2010). Players were provided a set of five starting models from I-TASSER (Zhang, 2008). Players then modified the structure to minimize the Rosetta energy function. 339 and 443 players participated in the D1 and D6 puzzles, respectively, over a two week period. A very diverse set of models was generated, which were clustered to within 2 Å RMSD. The 1000 top-scoring models were then used for molecular replacement with PHASER. A candidate model was chosen with high log-likelihood gain and translational function Z-score for further manual refinement (Figure S2). The D6 and D1 models were added sequentially to the VEGFR-1 D2-D5/VEGF-A model, followed by manual rebuilding in COOT (Emsley and Cowtan, 2004) and refinement in PHENIX (Adams et al., 2002) as for the homology models. NCS, secondary-structure and reference structure restraints were applied throughout the refinement process. For the final refined structure, Ramachandran values were: 82.2% favored, 14.4% allowed and 3.4% outlier.

The structure of the VEGFR-1 D2-7/VEGF-A complex was determined to 4.8 Å resolution by molecular replacement in PHASER using the D2-6 part of the VEGFR-1 D1-6/VEGF-A structure and the structure of VEGFR-2 D7 dimer (PDB ID: 3KVQ) as search models. The structure was improved by rigid-body refinement in PHENIX. A composite model of the full-length VEGFR-1 extracellular domain/VEGF-A complex was constructed in COOT by secondary-structure match (SSM) superposition of the D1-6 and D2-7 structures (RMSD of 1.76 Å for 1128 residues of the D2-6 region).

*Native-SAD experiments.* Highly redundant diffraction data were collected from a very large native crystal (700 μm) of the VEGFR-1 D1-6/VEGF-A complex at an X-ray energy of 6 keV (2.0663-Å wavelength) applying a recently described method (Weinert et al., 2015). Individual datasets of 360° were measured from multiple positions and orientations of the crystal giving 4320° in total with the anomalous signal extending to 4.5 Å. The final VEGFR-1 D1-6/VEGF-A model was refined against these long-wavelength data in PHENIX, which allowed for calculation of anomalous difference Fourier maps for sulfur. The anomalous peaks around all disulfide bridges in Ig-domains and in the VEGF ligand and additionally around several methionine sulfurs confirmed the accuracy of the model.

## AUTHOR CONTRIBUTIONS

ES and SMM produced, purified and crystallized proteins, performed electron microscopic and crystallographic experiments and wrote the manuscript, MA performed the biochemical analysis of receptor mutants, ES and KK performed SAXS experiments, TW assisted in diffraction data acquisition, structure phasing and validation, SB initiated and analyzed the FOLDIT molecular modeling, KNG assisted in electron microscopy data acquisition and analysis, GC was involved in structure refinement and manuscript preparation and KBH conceived and planned the experiments and wrote the manuscript.

## Supplemental Experimental Procedures

### Size exclusion chromatography coupled to multi-angle light scattering (SEC-MALS)

100 µl protein samples at a concentration of 1 mg/ml were injected onto a Superdex 200 HR 10/300 column for SEC-MALS determination (GE Healthcare). The column was equilibrated in 25 mM HEPES pH 7.5, 500 mM NaCl at 20°C, using an Agilent 1100 HPLC system. Light scattering and differential refractive index were recorded using a Wyatt miniDAWN Tristar detector and a Wyatt Optilab rRex detector, respectively. Wyatt Astra software was used to collect and process the data and to calculate the molar mass from a global fit of the light scattering signals from three detectors at different angles and the differential refractive index signal.

### Small angle X-ray solution scattering (SAXS)

SAXS data acquisition was performed at the X12SA-beamline (cSAXS) at the Swiss Light Source at the Paul Scherrer Institute in Villigen, Switzerland, details are given in Supplemental Experimental Procedures. The intensities of the scattered X-rays were recorded on a Pilatus 2M detector using a wavelength of λ = 1 Å. Data was collected in the scattering vector range of 0.008 Å^-1^-0.4 Å^-1^, where the length of the scattering vector is given as s = 4πsinθ/λ (2θ is the scattering angle). Silver behenate was used as a standard for calibration of the s-range10. Three concentrations were measured per protein sample using quartz capillaries with a diameter of 1 mm (Hilgenberg GmbH). To record background scattering, the protein buffer was measured in the same capillaries prior to protein data acquisition. Exposures of 0.5 s were taken at ten different spots along the capillary. The data was monitored for radiation damage and all frames showing no radiation damage were merged and averaged for further data processing. The collected SAXS data were integrated and radially averaged utilizing our own MATLAB-scripts (J. Missimer & K. Kisko, unpublished). Using PRIMUS from the ATSAS (Petoukhov et al., 2012) program package14, the background scattering was subtracted and data of different concentrations were merged. In order to check for proper protein folding of all measured samples Kratky-plots were calculated. The distance distribution function P(r) was calculated using GNOM and the program AUTOGNOM.

### Negative stain single particle electron microscopy (EM)

In order to stabilize the VEGFR-1 D1-7/VEGF-A complex for negative stain EM, samples were prepared using GraFix (ref. (Kastner et al., 2008)). Briefly, 150 pmol of the complex were applied onto a 5-20 % glycerol gradient containing 20 mM HEPES pH 7.5, 150 mM NaCl in the presence or absence of 0.2% glutaraldehyde. The samples were centrifuged for 18 hrs at 40’000 rpm in a SW60Ti rotor (Beckman-Coulter), which correlates to an average RCF value of 164’326 × g. The gradients were harvested from bottom to top with a peristaltic pump collecting 120 μl fractions. Protein containing fractions were detected using Roti^®^-Quant reagent (Carl Roth GmbH + Co. KG) for protein quantitation according to Bradford. In addition, protein containing fractions were analyzed by SDS-PAGE to verify the presence of the complex. Cross-linking reactions were quenched with the addition of 5x quenching buffer (20 mM HEPES pH 7.5, 150 mM NaCl, 0.4 M glycine). For negative stain EM, 5 μl of undiluted sample was adsorbed to a freshly glow discharged thin carbon film supported on a 200 mesh copper grid and incubated for 1 min. at room temperature. The excess sample was blotted away using filter paper. Three consecutive washes using 20 mM HEPES pH 7.5, 150 mM NaCl and one wash with 2% uranyl acetate was performed followed by 20 s staining with fresh 2% uranyl acetate. Negative stain EM data were collected on a CM-100 microscope (Philips) equipped with a Veleta 2k × 2k CCD camera (Olympus). The voltage used was 80 kV and the magnification was set to a nominal value of 130’000 x. From 126 micrographs 4883 particles were picked manually using XMIPP (Scheres et al., 2008). Inferior particles and particles with a z-score >3 were discarded. 4536 particles were boxed in 100 × 100 pixel boxes and subjected to alignment and classification into 24 2D class averages using maximum-likelihood target function (ML2D) implemented in XMIPP using 100 iterations (Scheres et al., 2005). After 49 iterations ML2D reached convergence.

### Isothermal titration calorimetry (ITC)

The measurements on VEGFR-1 were performed at 20°C in 25 mM HEPES pH 7.5, and 150 mM NaCl (ITC-buffer) using an iTC200 calorimeter (MicroCal^®^, GE Healthcare). The proteins were purified on a Superdex 200 10/300 column (GE Healthcare) equilibrated in ITC-buffer and dialyzed against ITC-buffer overnight at 4°C prior to analysis. The calorimeter cell contained VEGFR-1 D1-7 or VEGFR-1 D1-3 at concentrations ranging from 4.4 to 30 μM and the ligand VEGF-A121 was used in the syringe at concentrations ranging from 22.5 to 100 μM. All samples were equilibrated to the measurement temperature and degassed prior to ITC analysis. The following settings were applied: one initial injection of 1 μl followed by 15 injections of 2.6 μl at an injection speed of 1 μl/s with a data filter of 1 s and 300 s recovery time between each peak. The software Origin 7.0 (OriginLab^®^) was used for data analysis.

### Cell transfection and determination of receptor activity

NIH 3T3 cells were transfected using a modified peGFP-N1 vectors (Takara Bio Europe/Clontech) carrying either wt VEGFR-2, or a mutant T446E/K512A where residue K512 was replaced by A and T446 by E. In addition, a chimeric VEGFR (VEGFR-1/2) consisting of the wt or a mutant VEGFR-1 ECD (residues 1-752) and the intracellular domain of VEGFR-2 (residues 759-1356) was generated. The mutant used was E513K/K517A. All expression vectors were generated by Gibson assembly(Gibson et al., 2009). 8 µg DNA in 0.8 ml serum-free medium were mixed with 16 µl PEI (Polyethyleneimine, stock 1 mg/ml, Aldrich 408727, 25 kDa branched), incubated for 10 minutes at RT for the complex to form. The medium in the culture dishes was replaced with 1.6 ml DMEM containing 0.5% serum and the DNA/PEI complex was added for 5 hrs. Cells were incubated for 20 hrs after addition of 1 ml DME with 10% serum. The cells were serum-starved in DMEM supplemented with 1% bovine serum albumin (BSA) and stimulated with 1.5 nM VEGF-A for 10 min at 37°C. Cell lysates were prepared in lysis buffer (50 mM Tris pH 7.5, 100 mM NaCl, and 0.5% w/v Triton X-100) supplemented with protease inhibitor cocktail (Roche diagnostics, Risch-Rotkreuz Switzerland), phosphatase inhibitors (200 μM Na3VO4, 10 mM NaF, 10 mM sodium pyrophosphate, 30 mM p-nitrophenyl-phosphate, 80 mM glycerophosphate, and 20 μM phenylarsine oxide), and 10% glycerol. Cell lysates were diluted with Lämmli buffer (20 mM Tris pH 6.8, 5% SDS, 10% mercaptoethanol, 0.02% bromphenol blue) and the proteins separated on 8% SDS polyacrylamide gels and blotted to Polyvinylidene (PVDF, Thermofisher Scientific, Zug Switzerland) membranes. We used 55B11 (2479, Cell Signaling Technology Europe) and phospho-Y1175 (2478, Cell Signaling Technology Europe) for VEGFR-2 and VEGFR-1/2 detection. All primary antibodies were used at dilutions of 1:1’000, followed by secondary alkaline phosphatase-coupled antibodies (6440-04, Southern Biotech, Birmingham USA) at a dilution of 1:10’000. The blots were developed with AP chemiluminescent detection substrate (Life Technologies, Carlsbad USA) and the immunoblots were analyzed on a GE Healthcare ImageQuant RT ECL scanner and densitometrically quantified with ImageQuant TL software (GE Healthcare Europe GmbH, Glattbrugg, Switzerland).

## ACKNOWLEDGMENTS

K. B.-H. thanks Swiss National Science Foundation (grant 31003A-130463) and Oncosuisse (grant OC2 01200-08-2007) for continuous support of his research. S. B. was supported by the Intramural Research Program of the National Center for Biotechnology Information, National Library of Medicine, National Institutes of Health USA. We are also grateful to Thomas Schleier, Kate Thieltges and Julia Kostin for technical assistance, and the staff of the X06SA and X06DA beamlines of the Swiss Light Source for their support during data collection. We also thank the Foldit players and Firas Khatib for modeling D1 and D6 (https://fold.it/portal/info/credits).

## Supplemental Figures

**Figure S1.**
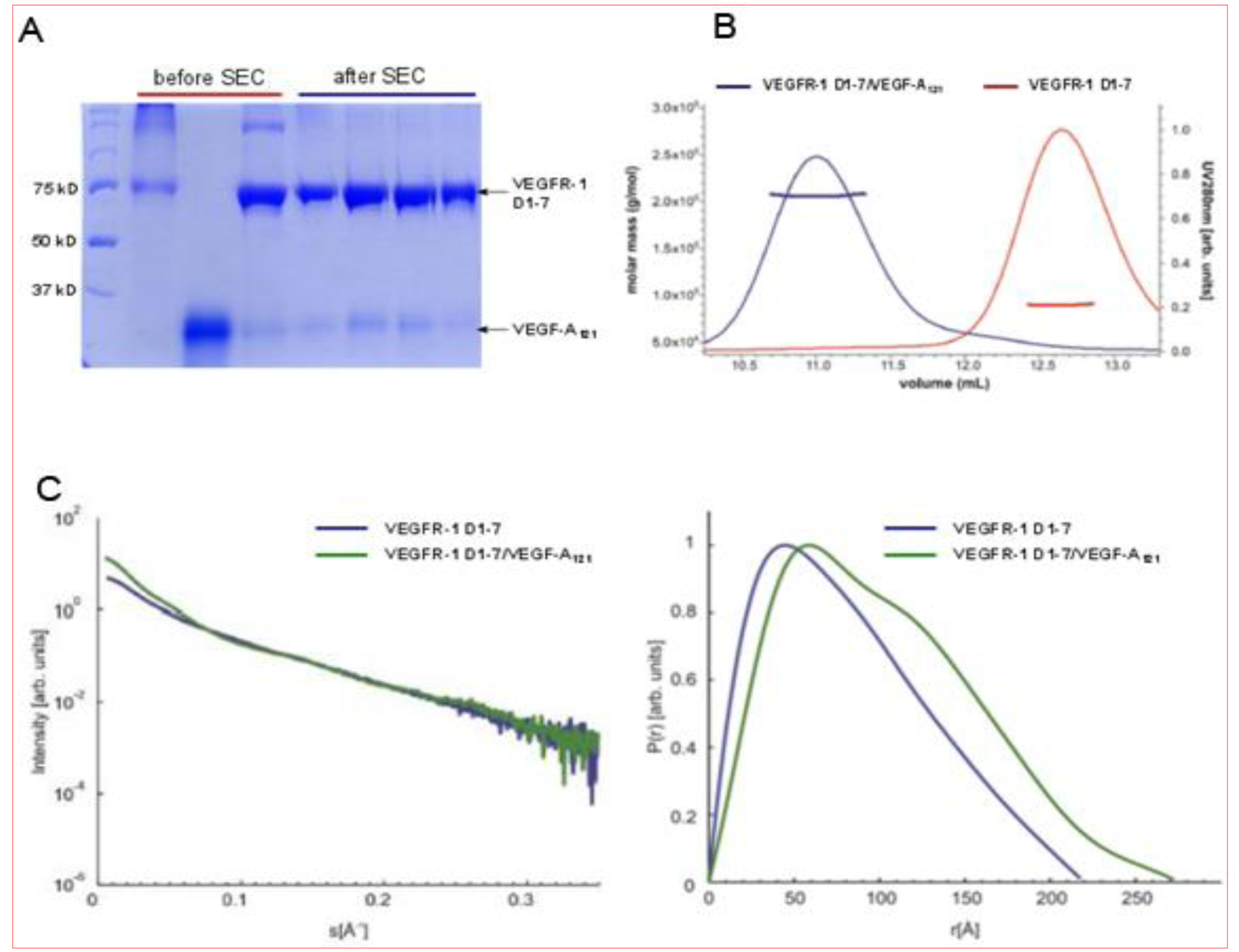
Figure S1 related to Figure 1 and 6. Purification and biophysical characterization of the VEGFR-1 ECD monomer and the VEGFR-1 ECD/VEGF-A complex in solution. **A**, SDS-PAGE analysis of purified VEGFR-1 ECD, VEGF-A_121_and the VEGFR-1 ECD/VEGF-A_121_complex. The preformed complex was purified by size exclusion chromatography (SEC) on Superdex 200 and used for further biophysical and structural analyses. **B**, Size exclusion coupled to multi-angle light scattering (SEC-MALS) of VEGFR-1 ECD alone and in complex with VEGF-A_121_. The UV-profiles are shown along with the mass distribution of each peak calculated from the MALS data. Experimentally determined Mr values were, for the VEGFR-1 ECD 89.7 kDa and for the VEGFR-1 ECD/VEGF-A_121_ complex 206.3 kDa (theoretical values 83.3 and 196.8 kDa, respectively). **C**, SAXS analysis of VEGFR-1 ECD and of the complex with VEGF-A_121_. Left panel: Scattering intensity as a function of scattering vectors. Right panel: GNOM distance distribution functions.

**Figure S2.**
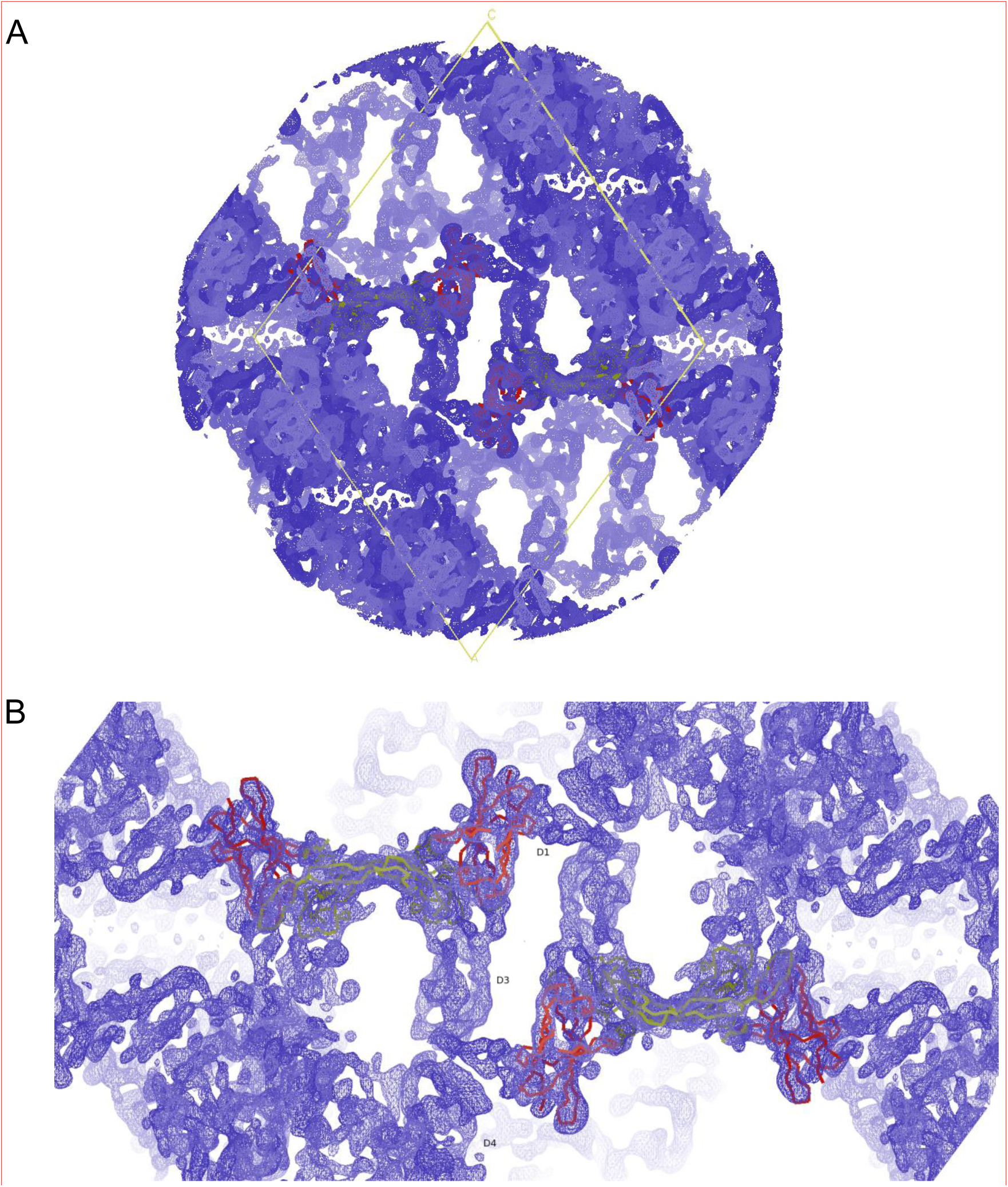
Figure S2, related to Figure 1. Quality of the experimental electron density map. Electron density map as obtained from MR-SAD phasing in PHASER and subsequent density modification in DM. The map is contoured at 1.5 σ and exhibits clear solvent-protein boundaries. The figure is centered on two VEGFR-1 D2/VEGF-A complexes positioned by molecular replacement and related by 2-fold crystallographic symmetry around y axes (VEGFR-1 D2 is colored red, VEGF-A is colored yellow). **A**, A view on the whole unit cell along the y axis. **B**, Zoom-in of panel a. The density corresponding to Ig-domains D1, D3 and D4 is indicated by labels.

**Figure S3.**
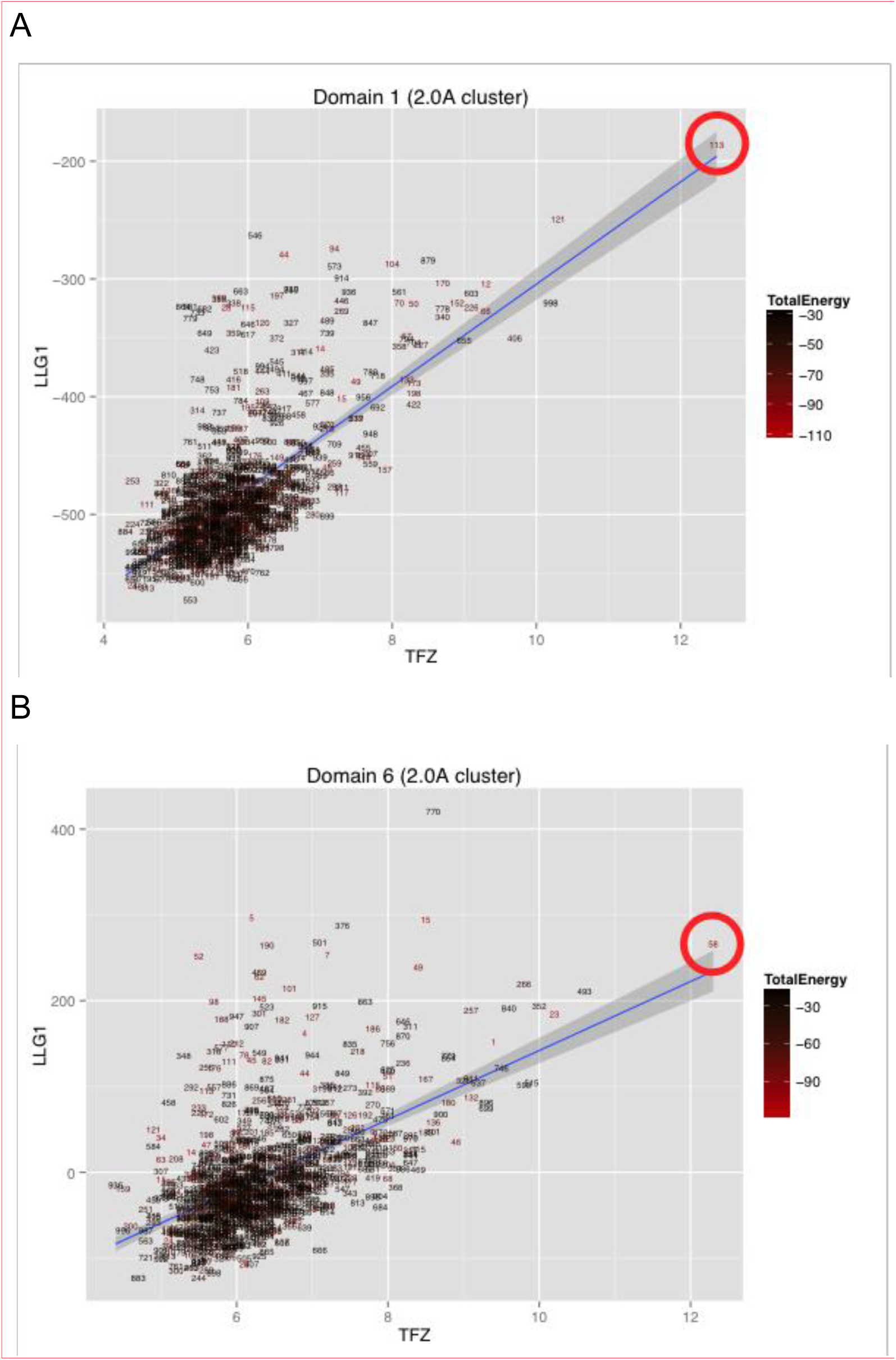
Figure S3 related to Figure 1. PHASER scores for molecular replacement of Foldit models for Ig-domain 1 (A) and Ig-domain 6 (B). Translation function z-scores (TFZ) and log-likelihood gains for the fitted domain (LLG1) were used to select a model for further use (red circle). Electron density fit only weakly correlated with Rosetta energy (Total Energy, color with labels giving ranked energy).

**Figure S4.**
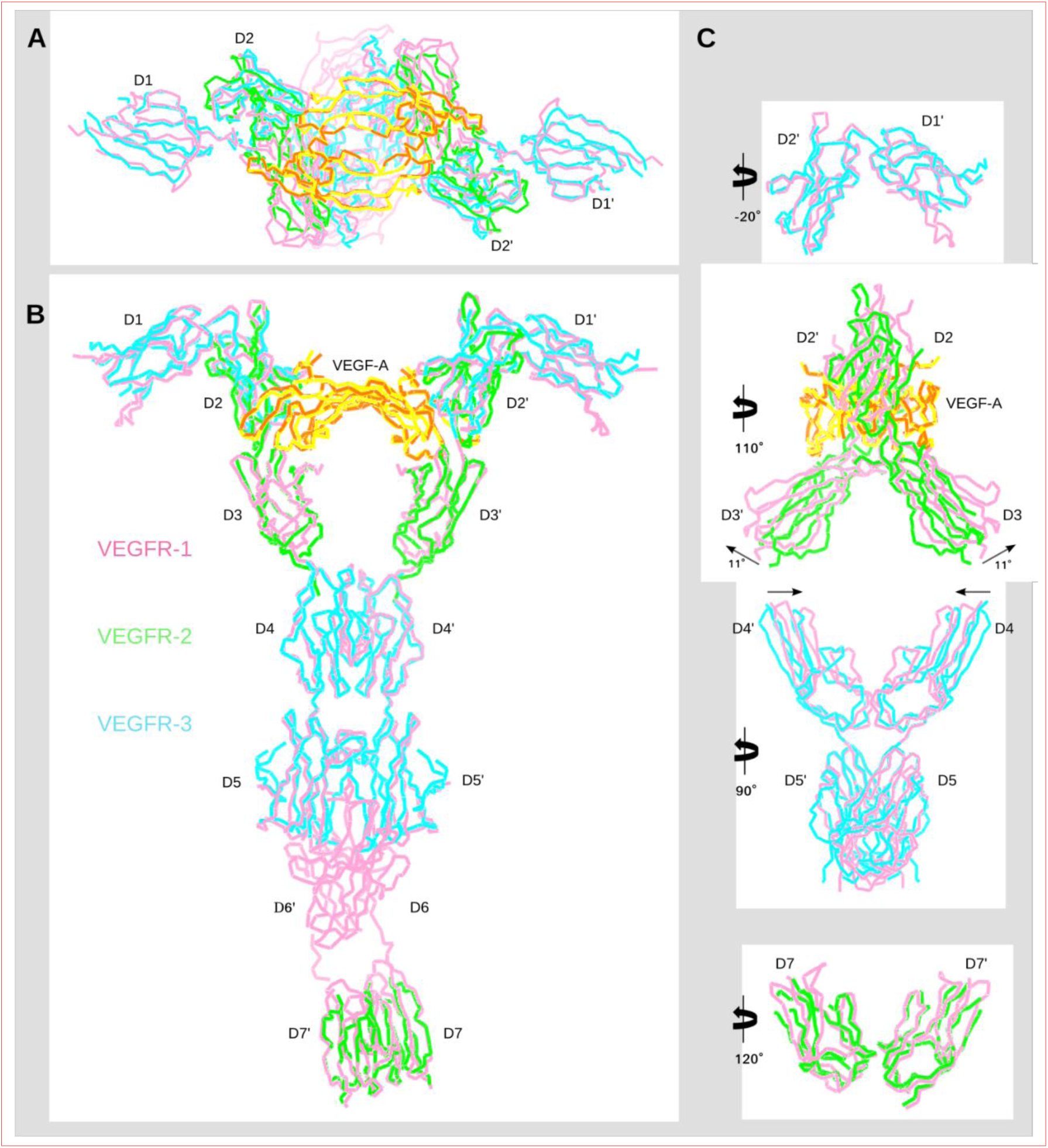
Figure S4 related to Figure 1. Comparison of the VEGFR-1 ECD/VEGF-A complex with known partial structures of VEGFR-2 and VEGFR-3. A, Top, and B, side views of the VEGFR-1 ECD / VEGF-A model (pink/yellow) with four known partial structures of VEGFR-2 (green) or VEGFR-3 (cyan) superimposed. From N-to C-terminus: VEGFR-3 D1-2 (PDB ID: 4BSK); VEGFR-2 D2-3 / VEGF-A complex (PDB ID: 3V2A) (green/orange); VEGFR-3 D4-5 dimer (PDB ID: 4BSJ); and VEGFR-2 D7 dimer (PDB ID: 3KVQ). Structural alignments were obtained by SSM superposition in Coot. C, The same four superpositions presented separately. Rotations relative to the orientation in B are designated.

**Figure S5.**
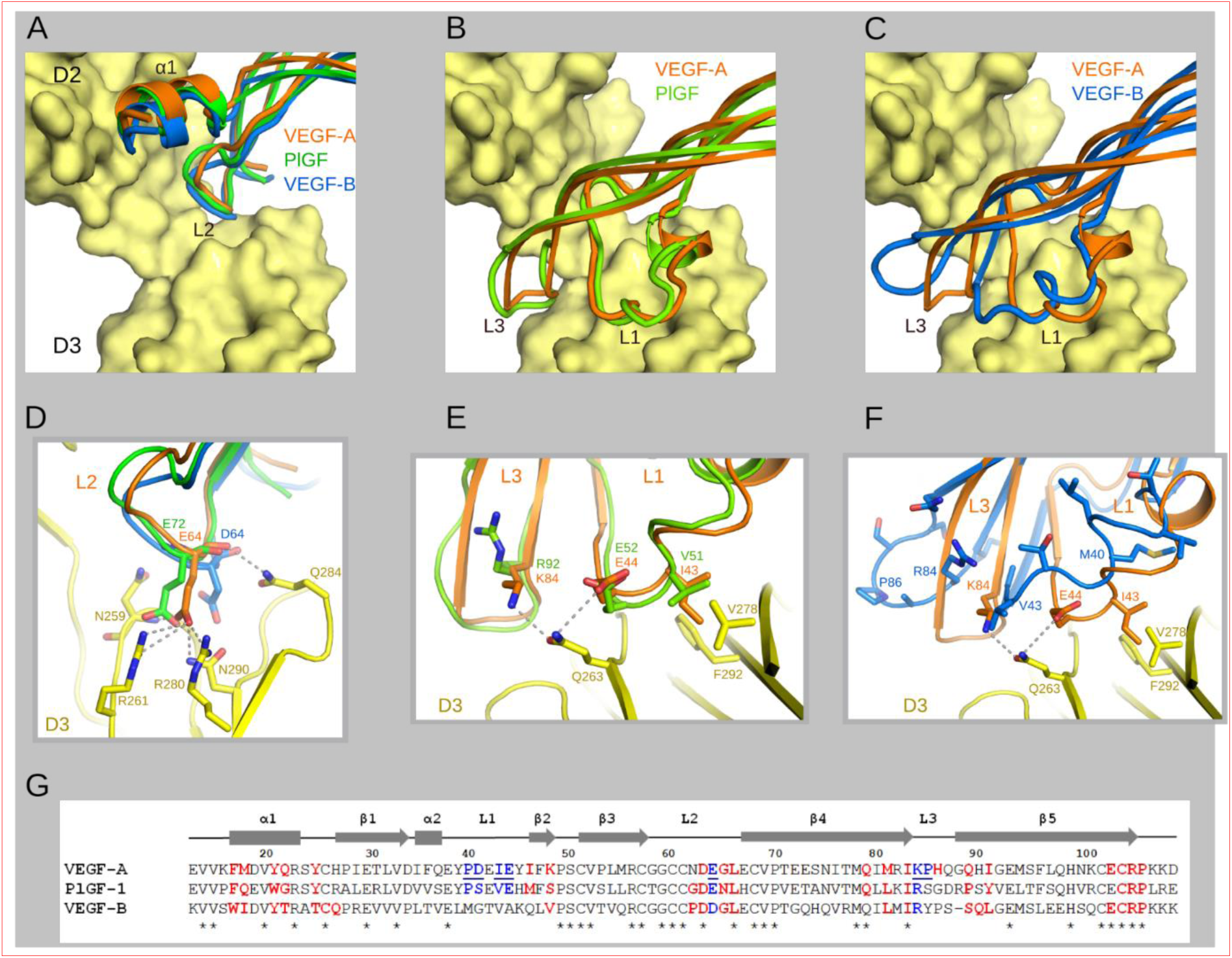
Related to Figure 4. Structural comparison of receptor binding epitopes in VEGF-A, PlGF and VEGF-B. A-C, Interaction of VEGF-A (orange) with VEGFR-1 D2-3 (yellow) as in Figure 4b. VEGFR-1 D2 complexes with PlGF (PDB ID: 1RV6) and VEGF-B (PDB ID: 2XAC) were superimposed onto D2 of the VEGFR-1 D1-6/VEGF-A complex structure. Parts of the ligands are shown: **A**, monomer A, helix α1 and loop 2; **B** and **C**, monomer B, loops 1 and 3. **D**, Interaction of E64 of VEGF-A with VEGFR-1 D3 and comparison with the corresponding residues in PlGF (E72) and VEGF-B (D64). Hydrogen bonds and salt bridges are shown as gray dashed lines. **E, F**, Interaction of VEGF-A L1 and L3 with VEGFR-1 D3 with the same loops of PlGF (**E**) and VEGF-B (**F**) superimposed. There is a high sequence and structure homology between L1 and L3 of VEGF-A and PlGF. L1 and L3 of VEGF-B have clearly different structural features. **G**, Structure-based sequence alignement of VEGF-A (13-109), PlGF (21-115) and VEGF-B (13-109). Residues that interact with VEGFR-1D2 are colored in red. VEGF-A residues that interact with VEGFR-1 D3 are colored in blue and underlined and the homologues residues in PlGF and VEGF-B that could play similar roles are colored in blue.

**Figure S6.**
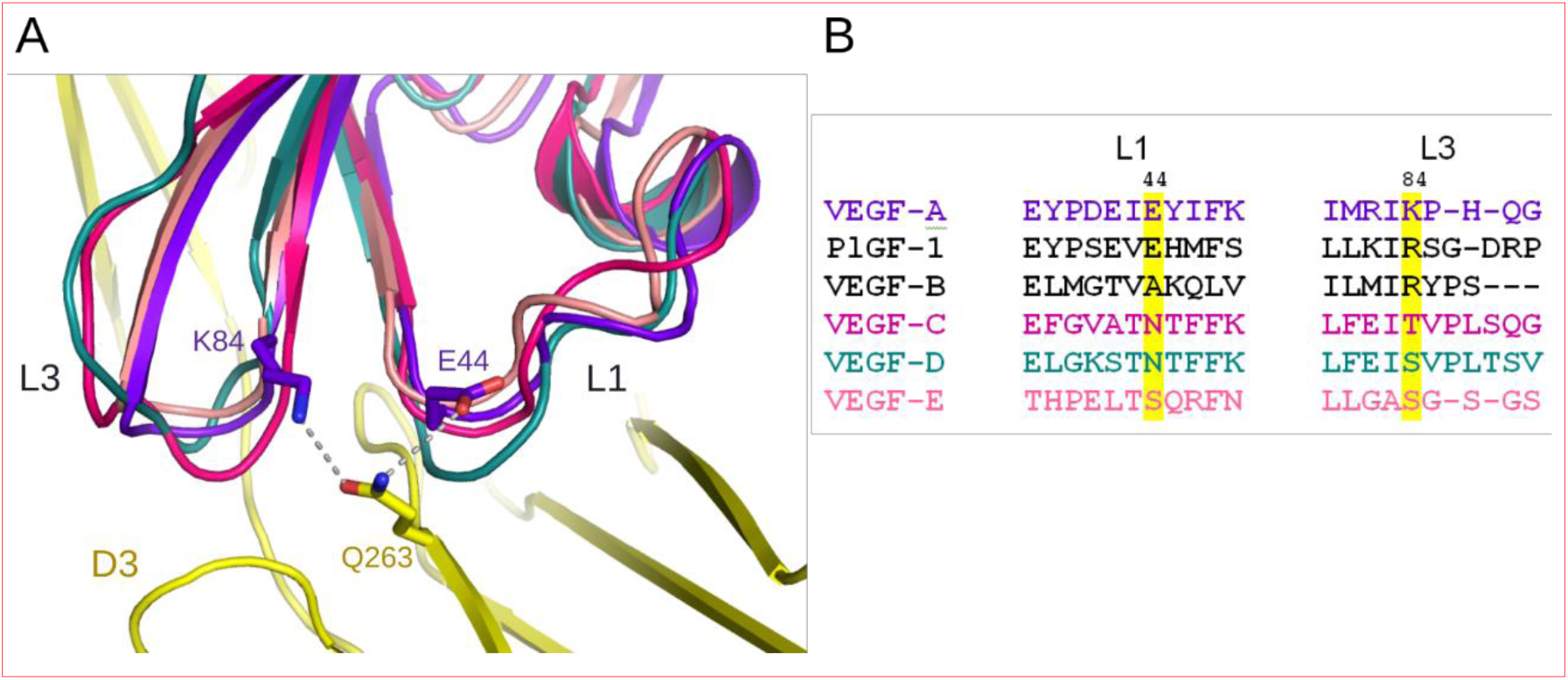
Figure S6 related to Figure 4. Structural comparison of the loops L1 and L3 of different VEGF family members. A,Interactions between the loops L1 and L3 of VEGF-A (purple) and VEGFR-1 D3 (yellow). The key interacting residues are shown as sticks and hydrogen bonds as gray dashed lines. VEGF-C (magenta, PDB ID: 2 × 1W), VEGF-D (deep teal, PDB ID: 2XV7) and VEGF-E (salmon, PDB ID: 3V6B) were superimposed onto our VEGFR-1 structure to compare L1 and L3 loops. **B**, Sequence alignement of the L1 and L3 regions of different VEGFs. VEGF-A residues E44 and K84 that interact with Q263 of VEGFR-1 and their counterparts in other VEGFs are highlighted in yellow.

**Figure S7.**
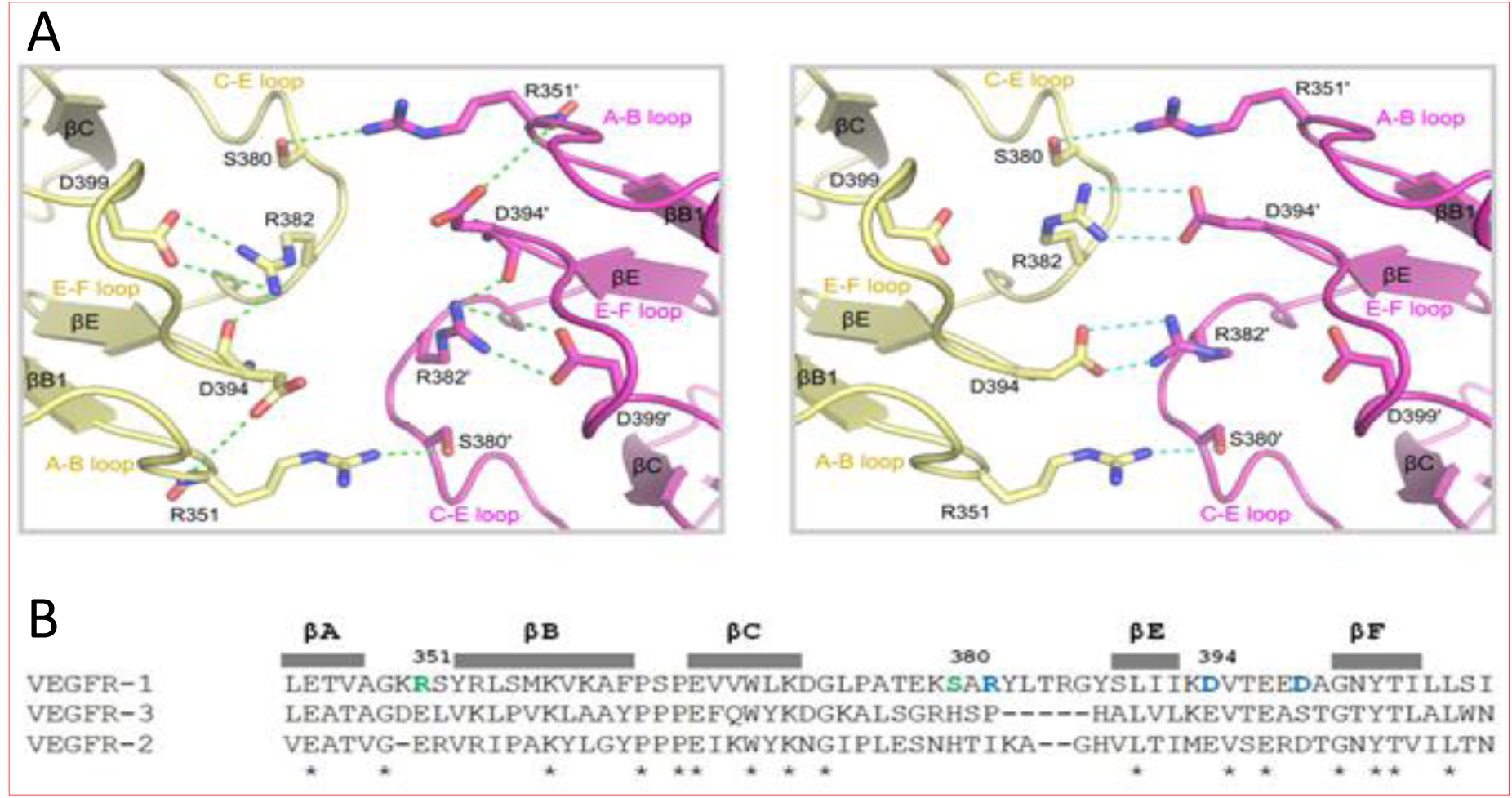
Figure S7 related to Figure 5. A closer view of the D4-D4 interface.A, Left panel–The interface as seen in the structure of the VEGFR-1 D1-6/VEGF-A complex with important residues shown as sticks and hydrogen bonds and salt bridges as green dashed lines. Right panel – Possible interface obtained by modification of side chain chi-angles for the residues R382 and D394 illustrates similarities with interactions observed between D4s of KIT (PDB ID: 2E9W) and D7s of VEGFR-2 (PDB ID: 3KVQ). **B**, Sequence alignment between D4 domains of three human VEGF receptors. R351 and S380 are colored in green, and the residues that might make additional contacts in blue.

